# PBMC Fixation and Processing for Chromium Single-Cell RNA Sequencing

**DOI:** 10.1101/315267

**Authors:** Jinguo Chen, Foo Cheung, Rongye Shi, Huizhi Zhou, Wenrui Wenrui, CHI Consortium, Julián Candia, Yuri Kotliarov, Katie R. Stagliano, John S. Tsang

## Abstract

**Background:** Interest in single-cell transcriptomic analysis is growing rapidly, especially for profiling rare or heterogeneous populations of cells. In almost all reported works investigators have used live cells, which introduces cell stress during preparation and hinders complex study designs. Recent studies have indicated that cells fixed by denaturing fixative can be used in single-cell sequencing, however they did not usually work with most types of primary cells including immune cells.

**Methods:** The methanol-fixation and new processing method was introduced to preserve human peripheral blood mononuclear cells (PBMCs) for single-cell RNA sequencing (scRNA-Seq) analysis on 10X Chromium platform.

**Results:** When methanol fixation protocol was broken up into three steps: fixation, storage and rehydration, we found that PBMC RNA was degraded during rehydration with PBS, not at cell fixation and up to three-month storage steps. Resuspension but not rehydration in 3X saline sodium citrate (SSC) buffer instead of PBS preserved PBMC RNA integrity and prevented RNA leakage. Diluted SSC buffer did not interfere with full-length cDNA synthesis. The methanol-fixed PBMCs resuspended in 3X SSC were successfully implemented into 10X Chromium standard scRNA-seq workflows with no elevated low quality cells and cell doublets. The fixation process did not alter the single-cell transcriptional profiles and gene expression levels. Major subpopulations classified by marker genes could be identified in fixed PBMCs at a similar proportion as in live PBMCs. This new fixation processing protocol also worked in several other fixed primary cell types and cell lines as in live ones.

**Conclusions:** We expect that the methanol-based cell fixation procedure presented here will allow better and more effective batching schemes for a complex single cell experimental design with primary cells or tissues.

## Background

The study of individual immune cells, the fundamental unit of immunity, has recently transformed from phenotypic analysis only to both phenotypic and transcriptomic analysis [1, 2]. This shift has been driven by the rapid development of multiple single-cell technologies in the last few years [3, 4]. Rather than studying population-averaged measurement, the modern single-cell RNA sequencing (scRNA-Seq) approaches have proved invaluable for identifying cell subtypes, especially rare cell populations; discovering highly variable genes contributing to cell-to-cell heterogeneity; and measuring individual cell responses to specific stimuli. Compared with the previously existing methods such as sorting-based microwell plates or microfluidics-based Fluidigm C1 [5], droplet-based techniques have enabled processing of tens of thousands of cells in a quick and unbiased way with trivial effect on cells [6]. Commercially available Chromium system manufactured by 10X Genomics greatly improves the cell capture efficiency and standardizes the protocol [7]. Hundreds to tens of thousands of cells are processed in under 7 minutes, with cell lysis beginning immediately after encapsulation into a droplet environment. It has emerged as the most widely used platform in the field of single-cell Sequencing.

The current scRNA-Seq protocols usually require using live cells. Molecular analysis of live cells, however, can be hindered by a variety of factors. Specifically, certain primary cell types, such as blood monocytes, rapidly undergo changes once isolated from whole blood. Fixation can stop cell stress/perturbation during the experiment. For complex experimental designs, development of preservation storage and successful resuscitation methods across a diverse number of cell types is essential for disconnecting time and location of sampling from subsequent single-cell sequencing experiments. Cell fixation can ease the logistic coordination. When multiple samples are analyzed, technical variation is commonly found in high throughput data [8]. Cell fixation may reduce batch and other confounder effects. In addition, samples acquired from individuals infected with highly infectious pathogens, such as HIV or HCV, are often restricted to facilities with biosafety containment. These samples must be fixed/killed prior to processing and analysis outside of an appropriate biosafety facility. Therefore, cell fixation would eliminate potential barriers to studying single-cell transcriptomes. Moreover, fixing cells provides a snapshot of cellular states at a given time point, i.e. samples can be analyzed at the same physiological state.

The ideal fixation method should be simple, efficient, and have little or no impact on the transcriptome (Table S1). Some recent studies have indicated that cells fixed by denaturing fixative can be used in single-cell sequencing [9, 10]. Alles et al have developed a simple methanol-based fixation protocol [9]. Cells are dehydrated with pre-chilled 80% methanol and then stored at −20°C or −80°C for up to several months. After rehydration in PBS, the fixed cells can be applied to subsequent profiling of single-cell transcriptomes by Drop-seq. The single-cell sequencing can be successfully performed with fixed cell lines such as HEK, 3T3, Hela, or fixed primary cells from some tissues such as *Drosophila* embryos and mouse brain. However, their protocol does not work in most primary cell types including lymphatic and immune relevant tissues such as peripheral mononuclear cells (PBMC), which are important targets of single-cell RNA-Seq. These cell types contain higher content of proteases and RNases than brain tissue (RNase Activity in Mouse Tissue, ThermoFisher TechNotes 12–3). Another issue not yet well addressed is whether there is RNA leakage or loss after cell fixation which could happen even if there is no RNA degradation [11]. In addition, single-cell analysis usually skips the RNA isolation step. If RNA leaks through the pores on the cell membrane into the suspension, the ambient (background cell-free) RNA concentration will go increase. When sequencing, these background reads cannot be related back to any specific cell.

To remedy these problems, we assessed these methanol-based fixation protocols [9, 10] and broke up the cell prep method into three sub-steps: 1) methanol fixation, 2) storage at −20°C, and 3) resuspension with PBS, to determine the steps at which the RNA degrades and loss occurs. We found that RNA from PBMCs was almost completely degraded during rehydration with PBS, not during cell fixation and storage. Resuspension in 3X saline sodium citrate (SSC) buffer instead of PBS protected PBMC RNA with RNA integrity number (RIN) greater than 8.0. We demonstrate that the methanol-fixed, SSC-resuspended PBMCs can be successfully implemented into the 10X Chromium standard scRNA-Seq workflows.

## Methods

### Single cell preparation

Human PBMCs were obtained from anonymous, healthy donors from the NIH Blood Bank. Cells were separated with LeucoSep tube filled with Ficoll-Paque-plus (GE Healthcare, Pittsburgh, PA) according to the manufacturer’s instruction. CD8+ cells were isolated from PBMC using Dynabeads^™^ CD8 Positive Isolation Kit (TermoFisher Scientific, Waltham, MA). Three other primary cells including <u>Human Lung Microvascular Endothelial Cells</u> (HMVEC-L), Human Bone Marrow Stromal Cells (BMSC) and mouse spleen cells were kindly provided by our collaborators. Culture cell lines, including KLM1, 293 T and MEF were harvested with trypsin-EDTA and single-cell suspension was prepared following 10X Genomics Single cell protocols: Cell preparation guide (CG00053, Rev C). Both PBMC and cell lines were washed twice to remove ambient RNA and finally resuspended in 1X PBS (calcium and magnesium free) containing 0.04% BSA. Cell concentration and viability were determined twice on a Guava^®^ easyCyte Single Sample Flow Cytometer (MilliporeSigma, Burlington, MA) using Guava^®^ ViaCount^®^ Assay. Cells with viability of greater than 90% were used and kept on ice for fixation and single cell RNA-Seq analysis.

### Cell fixation and post-fixation processing for Chromium^™^ scRNA-Seq

Methanol fixation was adapted from Alles et al [9] and Cao et al [10]. Between ~0.1 × 10^6^ (limited sample) and 1.0 × 10^6^ cells (general sample) in 1 volume (50 μl or 200 μl) of cold PBS-0.04%BSA were fixed with 4 volumes (200 μl or 800 μl) of 100% methanol (CH3OH, pre-chilled to −20°C). To avoid cell clumping, methanol was added dropwise, while gently stirring the cell suspension with micropipette tip. The cells were fixed at −20°C for 30 minutes and then stored at −20°C or −80°C until use (for up to 3 months). In order to check RNA quality and quantity, the cells were collected right after fixation (no wash step) or after storage and lysed in QIAzol. Total RNA was isolated and purified with miRNeasy kit (Qiagen, Hilden, Germany).

For resuspension, cells were removed from −20°C or −80°C and kept at 4°C throughout the procedure. Fixed cells were pelleted at 1000 g for 5 min. Methanol-PBS solution was completely removed. The cells were then resuspended in a small volume of cold SSC cocktail (3 X SSC-0.04%BSA-1% SUPERase•In^™^ −40 mM DTT) to keep a density of about 2000 cells/ μl. 20XSSC was purchased from KD Medical, Columbia, MD, SUPERase•In^™^ from Ambion, Austin, TX and dithiothreitol (DTT) from Invitrogen, Grand Island, NY.

The cell suspension was recounted before gel bead-in-emulsion (GEM) generation. For control of RNA quality after resuspension, cells were resuspended in the above SSC or PBS at 4°C for 30 min. A 50 µl cell suspension aliquot was mixed with 700 μl of QIAzol followed by total RNA isolation as above. Assessment of RNA quality was performed using the Agilent 2100 Bioanalyzer (Agilent Technologies, Santa Clara, CA).

### Full-length ds-cDNA synthesis using template switching technology

SMART-Seq v4 Ultra-low Input RNA kit for Sequencing (Takara, Mountain View, CA) was used to generate full-length ds-cDNA from total RNA according to the manufacturer’s instruction. This kit incorporates the Clontech’s SMART^®^ (Switching Mechanism at 5’ End of RNA Template) technology. Briefly, the 8 ng of control RNA was used as the template. First-strand cDNA synthesis from control RNA was primed by the 3’ SMART-Seq CDS Primer II A and used the SMART-Seq v4 Oligonucleotide for template switching at the 5’ end of the transcript. The full-length ds-cDNA from the SMART sequences was amplified by Long Distance PCR. PCR-amplified cDNA was validated using Agilent’s High Sensitivity DNA Kit. Successful cDNA synthesis and amplification should yield no product in the negative control, and a distinct peak spanning 400 bp to 10,000 bp, peaked at ~2,500 bp for the positive control RNA sample, yielding approximately 3.4–17 ng of cDNA as described in the manual.

### Single-cell encapsulation, library preparation and sequencing

Droplet-based single-cell partitioning and single-cell RNA-Seq libraries were generated using the Chromium Single-Cell 3’ Reagent v2 Kit (10X Genomics, Pleasanton, CA) as per the manufacturer’s protocol based on the 10X GemCode proprietary technology [7]. Briefly, a small volume (<4 μl) of single-cell suspension at a density of some 2000 cells/μl was mixed with RT-PCR master mix and immediately loaded together with Single-Cell 3’ Gel Beads and Partitioning Oil into a Single-Cell 3’ Chip. The Gel Beads were coated with unique primers bearing 10X cell barcodes, unique molecular identifiers (UMI) and poly(dT) sequences. The chip was then loaded onto a Chromium Controller (10× Genomics) for single-cell GEM generation and barcoding. RNA transcripts from single cells were reverse-transcribed within droplets to generate barcoded full length cDNA using Clontech SMART technology. After emulsion disruption, cDNA molecules from one sample were pooled and pre-amplified. Finally, amplified cDNAs were fragmented, and adapter and sample indices were incorporated into finished libraries which were compatible with Illumine next-generation short-read sequencing. The final libraries were quantified by real – time quantitative PCR and calibrated with an in-house control sequencing library. The size profiles of the pre-amplified cDNA and sequencing libraries were examined by Agilent Bioanalyzer 2100 using a High Sensitivity DNA chip (Agilent).

Two indexed libraries were equimolarly pooled and sequenced on Illumina NextSeq 500 system using the NextSeq 500 High Output v2 Kit (Illumina, San Diego, CA) with a customized paired end, dual indexing (26/8/0/98-bp) format according to the recommendation by 10X Genomics. Using proper cluster density, a coverage around 250 M reads per sample (3000 ~ 5000 cells) was obtained corresponding to at least 50,000 reads/cell.

### scRNA-Seq data preprocessing, alignment, gene quantification and QA/QC

The sequencing data was analyzed using the Cell Ranger Pipeline (version 2.0.1) to perform quality control, sample demultiplexing, barcode processing, alignment and single-cell 3’ gene counting. Samples were demultiplexed with *bcl2fastq* v2.19.1.403 based on the 8-bp sample index, 10-bp UMI tags, and the 16-bp GemCode barcode. The 98-bp-long read 2 containing the cDNA sequence was aligned using STAR against the GRCh38 human reference transcriptome. UMI quantification, GemCode, and cell barcodes filtering based on error detection by Hamming distance were performed as described by Zheng et al [7]. Only confidently mapped, non-PCR duplicates with valid barcodes and UMIs were used to form an unfiltered data matrix. The barcodes with total UMI counts exceeding *10%* of the 99th percentile of the expected recovery cells (default=3000) were considered to contain cells and selected to produce a filtered gene-barcode matrix for further analysis. “Genes and transcripts (UMI counts) per cell” was used to compare the sensitivity of scRNA-Seq before and after cell fixation. “Fraction Reads in Cells” was determined by the fraction of cell-barcoded, confidently mapped reads with cell-associated barcodes to check the background of cell-free (ambient) RNA in cell suspension.

Single-cell RNA-Seq data QA/QC was also run on Partek Flow single cell module (Build version: 6.0.17.1206), and hg38_ensembl_release90_v2 was used for gene/feature annotation. Any PBMC with more than 7% of mitochondrial UMI counts was considered to be a low-quality cell [12]. PBMC GEMs with greater than 2500 genes expressed or CD8 GEMs with more than 2000 detected genes were checked in order to determine the rate of doublets. Any gene detected in less than three cells or a cell with less than 200 genes detected was excluded for downstream data analysis.

### Normalization and correlation of gene expression levels

Each individual sample was normalized separately by cell RNA content as default setting in “cellranger count” pipeline. Only genes that were detected in at least three cells were included for the correlation and comparison, which used the mean of each gene expression across all cells. Fresh and fixed paired samples were also analyzed together with “cellranger aggr”. It normalizes multiple runs to the same sequencing depth (default=mapped) and then re-computes and produces a new single gene-barcode matrix containing all the data for correlation analysis.

### PCA and tSNE analysis for cell clustering and classification, and data visualization

The Cell Ranger count and aggr pipelines were used to run secondary analysis. Before clustering the cells, Principal Component Analysis (PCA) was run on the normalized, log-transformed, centered and scaled gene-barcode matrix to reduce the number of feature (gene) dimensions. The pipeline adopts a python implementation of IRLBA algorithm. This produced a projection of each cell onto the first N principal components (default N=10). It did not filter out any “low-quality” genes and cells as described above and previously [12] and used by Seurat package before PCA analysis. After running PCA, t-distributed Stochastic Neighbor Embedding (t-SNE) was run to visualize cells in a 2-D space. Clustering is then run to group cells together that have similar expression profiles, based on their projection into PCA space. Two clustering methods were performed: graph-based and k-means. Cell Ranger also produced a table indicating which genes were differentially expressed in each cluster relative to all other clusters. Classification of PBMCs was inferred from the annotation of cluster-specific genes, and was based on expression of some well-known markers of immune cell types (marker-based classification, Table S2). Loupe^™^ Cell Browser (v2.0) was used to view the entire dataset and interactively find significant genes, cell types, and substructure within cell clusters.

## Results

### RNA integrity was lost during rehydration with PBS, not during cell fixation and storage

Methanol is one of the most commonly used denaturing and precipitating fixatives for nucleic acid [10, 13, 14] and chromatin study [15]. It dehydrates cells/tissues, causing proteins and nucleic acids to denature and precipitate *in situ*. The complete removal of the fixative, even trace amounts, from the tissue or cell suspension must be carried out because it may impede subsequent processes or reactions. The cells are first pelleted and then washed with and resuspended in PBS, which is the most commonly used buffer. However this fixation and processing method causes RNA degradation [9, 16] and loss [11] in many types of cells, especially primary cells such as PBMC. To explore when RNA loses its integrity, we broke up the methanol fixation and processing procedure into three sub-steps: 80% methanol fixation, storage at −20°C and rehydration (resuspension) with PBS. At the end of each step, total RNA was checked with Bioanalyzer. As shown in Figure 1, RNA within PBMC was kept almost intact after methanol fixation and storage at −20°C for up to three months. High quality, intact RNA with RIN>8 could also be extracted from fixed KLM1 cells after storage in 80% methanol for two months. In addition, RNA content per cell did not change significantly during fixation and storage. However, RNA from PBMC resuspension in PBS after two-round washes had undergone extensive degradation (Figure. 1b), and completely degraded in 30 min. RNA quality from fixed KLM1 and 293 T cell lines was also compromised with RIN<8 after 30 min of incubation (Figure S1b).

**Figure 1.**
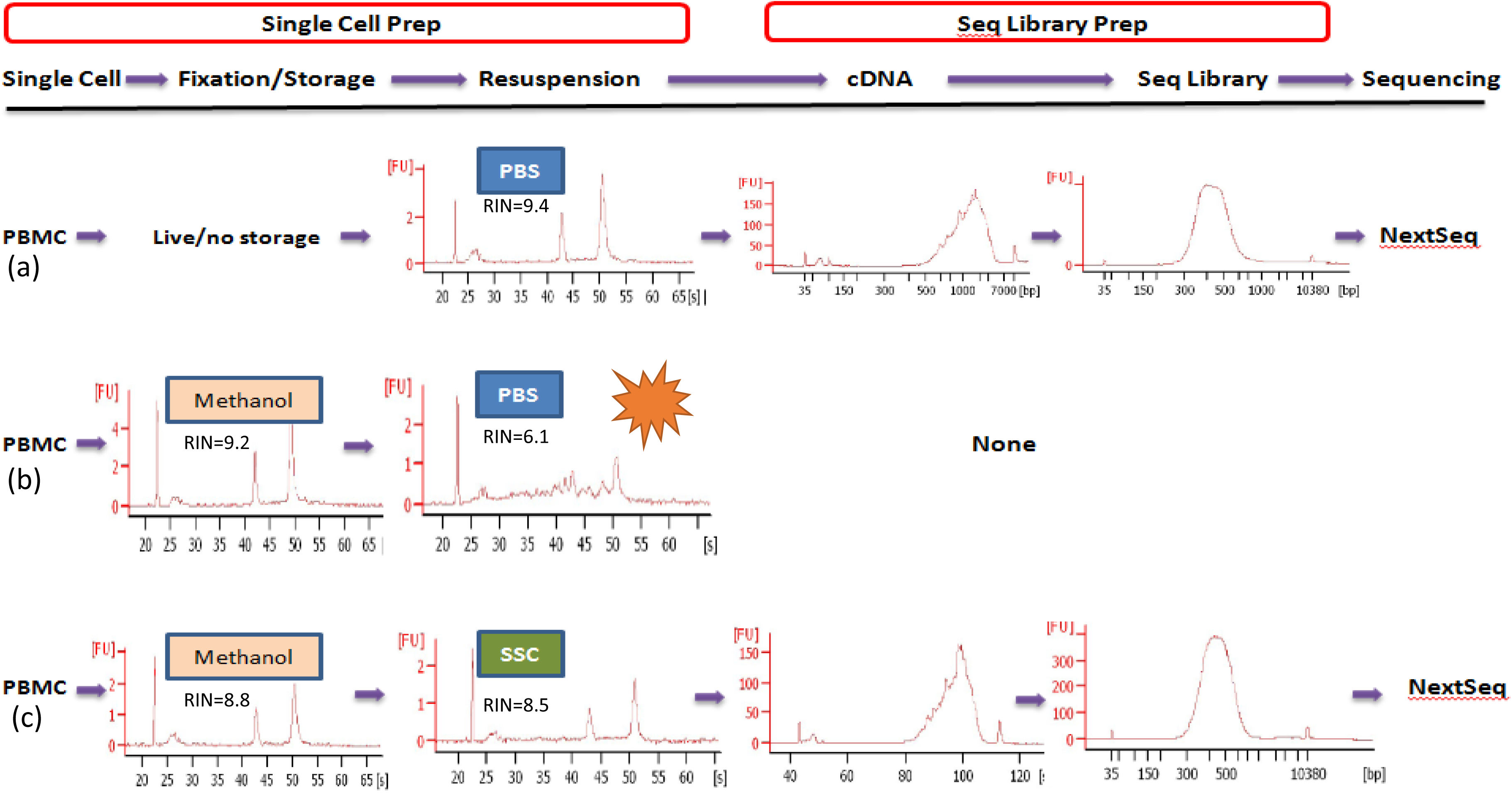
Experimental workflow of cell preparation and single-cell RNA-Seq. Donor PBMCs that were either live (**a**) or fixed (**b, c**) were analyzed their whole transcriptome at a single cell resolution. PBMCs in cold PBS (**b, c**) were fixed by adding 4 volumes of chilled methanol dropwise and then stored at −20°C or −80°C for up to three months. Right before scRNA-Seq, the PBMCs were resuspended in PBS (**b**) or SSC (saline sodium citrate, **c**). At the end of each step, the RNA quality was examined. **b**. Fixed PBMC RNA was degraded during rehydration with PBS and could not generate sequencing library. **c**. Fixed PBMCs resuspended in 3X SSC preserved RNA integrity with RIN>8.0 (confirmed in four different donors’ PBMCs) and were successfully implemented into 10X Chromium standard scRNA-Seq workflows as live PBMCs (**a**).

### Resuspension in 3x SSC buffer preserved PBMC RNA integrity

In order to find an appropriate resuspension solution, several RNA stabilization/preservation reagents or buffers were tested, including RNAlater and RNAProtect, and finally 3x or 5x SSC proved to be a good medium for the conservation of the fixed PBMCs and prevention of cellular RNA degradation. As shown in Figure S1a, RNA remained intact with RIN >8.0 when methanol-fixed cells were resuspended in 3x SSC or higher for 30 min. In addition, small RNAs such as 5S RNA were still retained in total RNA product. It is reported that smaller DNA fragments might be lost at concentrations of 10X SSC or less during membrane transfer and hybridization. In contrast, even high concentration (5%) of RNasin (Premega) or SUPERase•In^™^, a non-DTT-dependent formulation which offers broader protection against RNase, alone did not prevent fixed PBMC RNA from degradation. We also did not find protective effect of 40 mM DTT alone on PBMC RNA, which previously reported as working in three cell lines [17]. RNase inhibitor and DTT are the common components in reverse transcription reaction. They were thus added to the SSC suspension cocktail. BSA was also included in the SSC cocktail because fixed cells are sticky. BSA can block their nonspecific binding to tube. The protective effect of this SSC cocktail was confirmed in other three donors’ fixed PBMCs and one of isolated CD8+ T cells. It was also verified in three other primary cell types, including HMVEC-L, BMSC and mouse spleen cells and three cell lines such as KLM1, 293 T and MEF (Fig. S1b). Because of the high density of 3x SSC, cells in SSC suspension could not be pelleted before RNA isolation. At this point we could not know whether there was RNA leakage from cytoplasm to the medium (ambient RNA) after fixation and resuspension.

### Diluted SSC buffer did not have major impact on reverse transcription

SMART-Seq v4 Ultra-low Input RNA kit for Sequencing was used to investigate the possible inhibitory effect of SSC buffer on reverse transcription because it adapts the same chemistry to generate full-length ds-cDNA as Chromium Single Cell 3’ Reagent v2 Kit [7]. Different volumes of SSC were added to RT reaction solution with the control RNA sample provided in the kit. As shown in Figure 2a, 0.5X SSC did significantly suppress the cDNA synthesis, however when SSC was lowered to 0.125X, there was no inhibitory effect. The cDNA showed similar size distribution (600bp~9300bp), peak (~2100bp) and yield (8~10 ng) as those of control sample. Thus, this final concentration was used in the subsequent 10X reverse transcription reaction. It will not limit the number of cells loaded into the 10X chip if the cell concentration is high.

**Figure 2.**
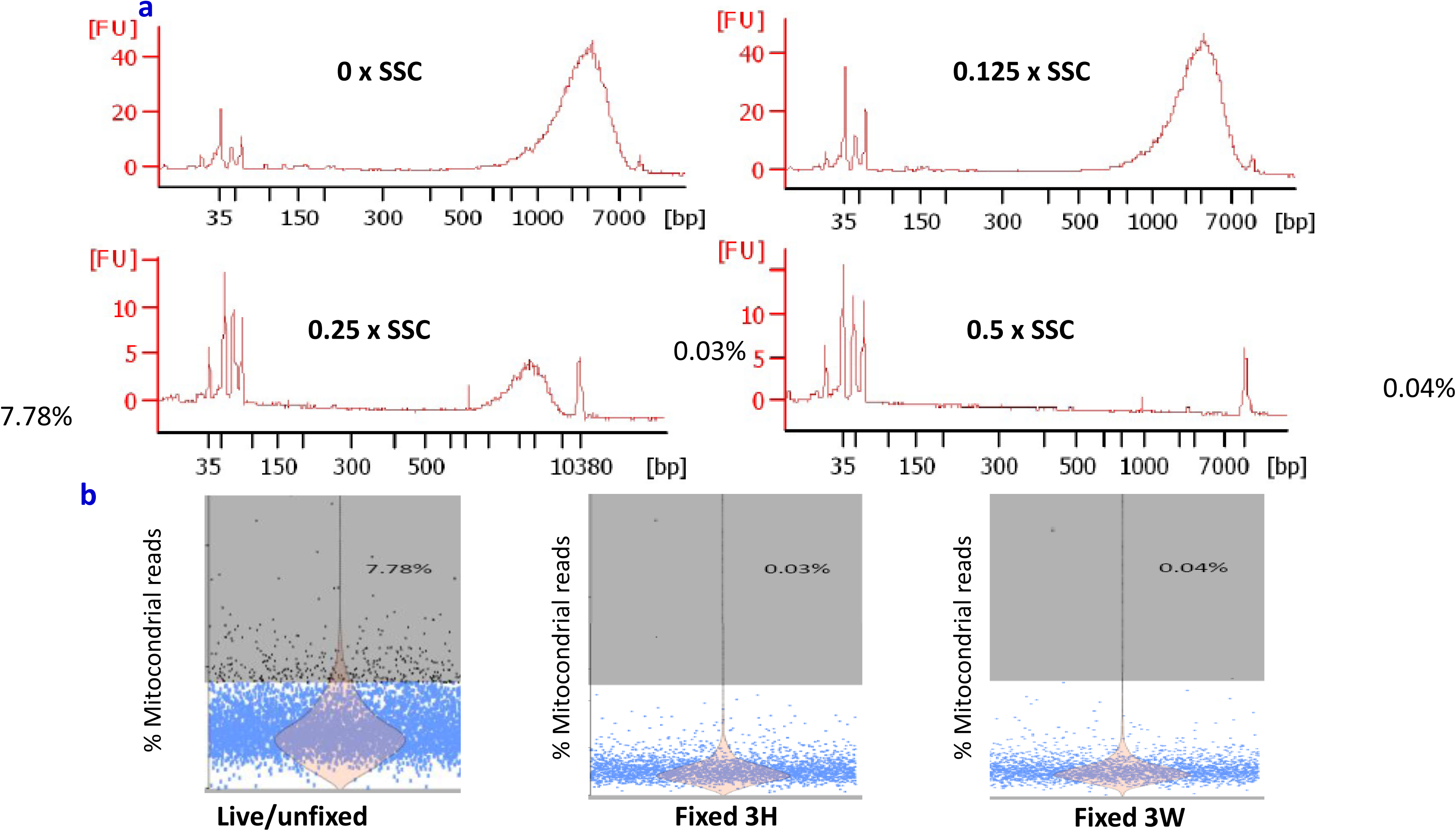
Diluted SSC neither interfered with full-length cDNA conversion nor produced low quality cells. **a**. When SSC was diluted to 0.125X, there was no inhibitory effect on SMART technology-based reverse transcription. The cDNA showed similar size distribution, peak and yield as that from control sample. Thus this final concentration was used in the subsequential scRNA-Seq. It will not limit the number of cells loaded into the 10X chip if the cell concentration is high. **b**. The percentage of 37 mitochondrial gene reads was calculated. High percentage (7% or higher highlighted in grey color) means cell suffered strong stress, and are broken or damaged to some degrees. Fixation did not increase these “low quality cells”.

### Low-quality cells and cell doublet rate were not elevated after fixation processing

An increase in the proportion of transcripts from mitochondrial genes is believed to indicate low-quality cells that are broken or damaged to some degrees [12]. We thus investigated if fixation processing resulted in more “low quality cell”. The percentage of 37 mitochondrial gene reads was calculated in each cell. High percentage (7% or higher) means cell suffered strong stress, leading to loss/leakage of cytoplasmic RNA, while mitochondrial located mRNA transcripts are protected by two layers of mitochondrial membranes. The proportion of mitochondrial mRNA had elevated in 7.78% of live PBMC sample after single-cell preparation (Figure 2b); however, this proportion went down to less than 1% in fixed cells. Thus, fixation processing did not seem to cause a rise in low-quality cells. In contrast, fixation prevented the PBMC from further stress/perturbation during prolonged cell manipulation.

Fixed cells are sticky. In order to assess whether fixed PBMCs easily aggregate to form doublets or multiplets, the GEMs with high number of detected genes were examined. These populations usually contain more than one cell. Partek Flow QC data indicated this ratio was kept low after fixation (Figure S2). In addition, fixation did not induce a microscopically detectable increase in cell aggregates. Methanol-fixed PBMCs remained visible as single, intact round cells with sizes similar to those of live ones.

### Fixation processing preserved single-cell RNA profiling and gene expression levels

To determine if methanol-fixed, SSC-conserved PBMCs can be applied to droplet-based 10X Chromium scRNA-Seq, we fixed two vials of fresh PBMCs from donor DTM-X, stored at −20°C for 3 hours (3 H) or 3 weeks (3 W), and constructed the scRNA-Seq libraries from these fixed cells resuspended in a small volume of the SSC cocktail described previously. The sequencing matrix was shown in Table S3. The cDNA and finally libraries in Bioanalyzer traces appeared indistinguishable between fixed and live samples (Figure 1). The sequence reads from three datasets (one live and two fixed) had similar alignment percentage to reference transcriptome. The medium genes and UMIs detected per fixed PBMC (Figure 3a) showed somewhat lower than those per fresh PBMC. The drop rate was about 20% in Donor X PBMC, Donor Y PBMC and CD4+ Cells (Table S3), indicating a consistent conversion efficiency of the system. The number of detected genes was still much higher than that reported with version 1 reagent by 10X company [7]. In addition, their average gene expression levels were highly correlated (Pearson’s correlation test, r= 0.95–0.97, Figure 3b), especially between two fixed PBMC samples (r=0.98) from one donor fixed at the same time but preserved for different duration and sequenced separately, demonstrating this new fixation method is quite reproducible. In theory, there is no biologically up- or down-regulation of gene expression after methanol fixation. We did find that only 2 or 15 gene expression levels increased with two or more fold change after fixation for 3 hours or 3 weeks respectively. They are more likely due to technical not biological variations.

**Figure 3.**
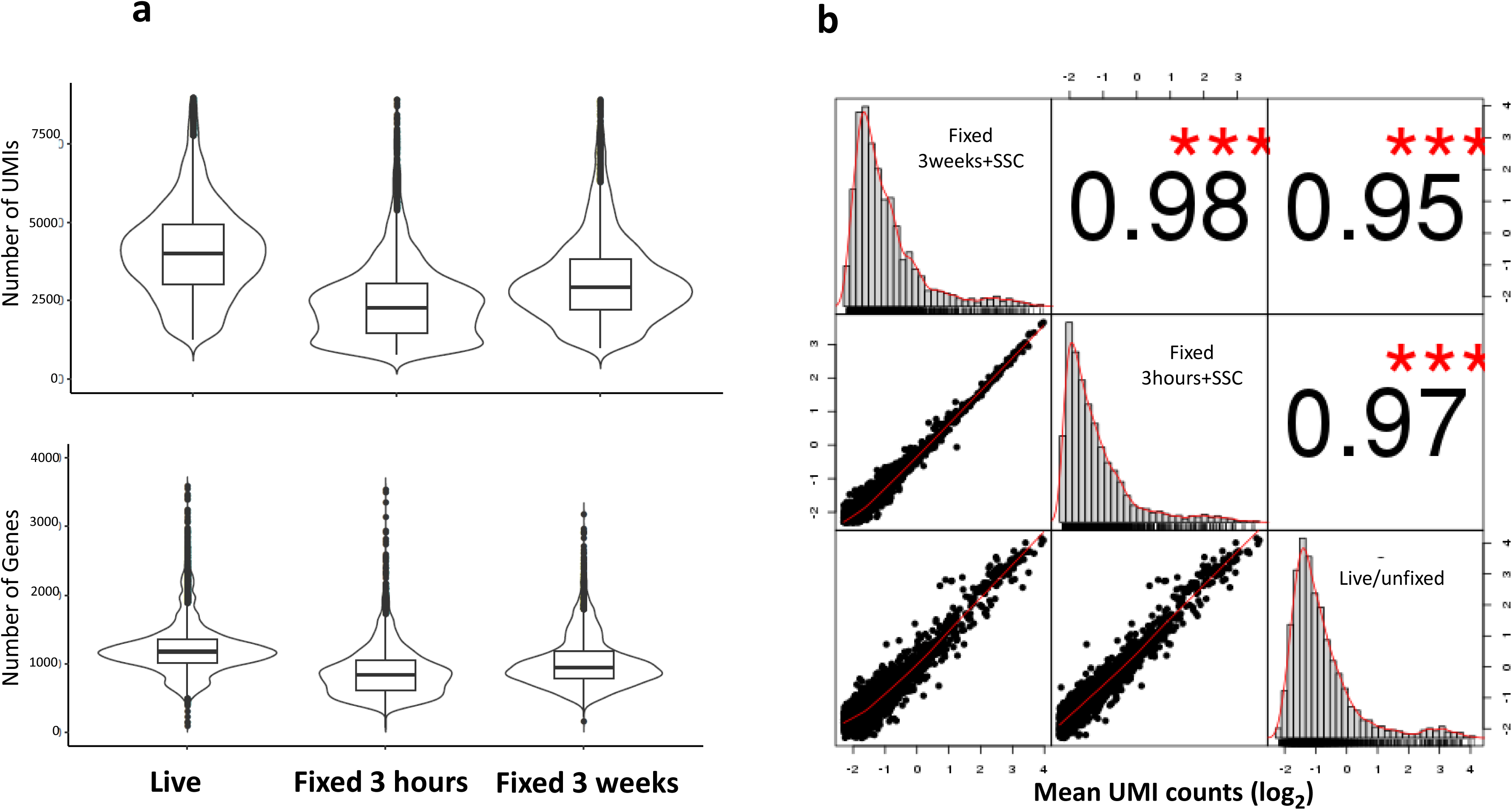
Fixation processing preserves single-cell RNA profiling and gene expression levels. **a**. Fixation did not affect the sensitivity of the single cell analysis. The UMI and genes detected per cell were slightly lower after fixation. The drop rate is about 20% in Donor X PBMC, Donor Y PBMC and CD4+ Cells, indicating a consistent conversion efficiency of the system. The number of detected genes was still much higher than that reported with version 1 reagent by 10X company. **b**. Gene expression levels (mean UMI/gene/across all cells) showed a high similarity among live and fixed cells, especially between two fixed PBMC samples fixed at the same time but preserved for different duration (3 H and 3 W) and sequenced separately, indicating our fixation method is highly reproducible. Only 2 and 15 gene expression levels increased with two or more fold changes after fixation for 3 hours and 3 weeks respectively. *** p<0.001

When methanol-fixed KLM1 was resuspended in PBS, their RNA was partially degraded (Figure 4a) and its “fraction reads in cells” was only 53.8%, much lower than that from live sample (Figure 4b). The genes and UMI counts detected also dropped by 20% and 30% respectively (Table S3), which also happened in other reported cell lines [9]. In contrast, KLM1 samples resuspended in SSC cocktail had much higher percent UMI counts associated with cell barcodes, indicating low ambient cell-free RNA. The genes and UMI counts detected were almost the same as those of live KLM1 cells (Figure 4c). In summary, SSC not only deterred RNA from degradation but also prevented the cytoplasmic RNA leakage after the fixative was removed.

**Figure 4.**
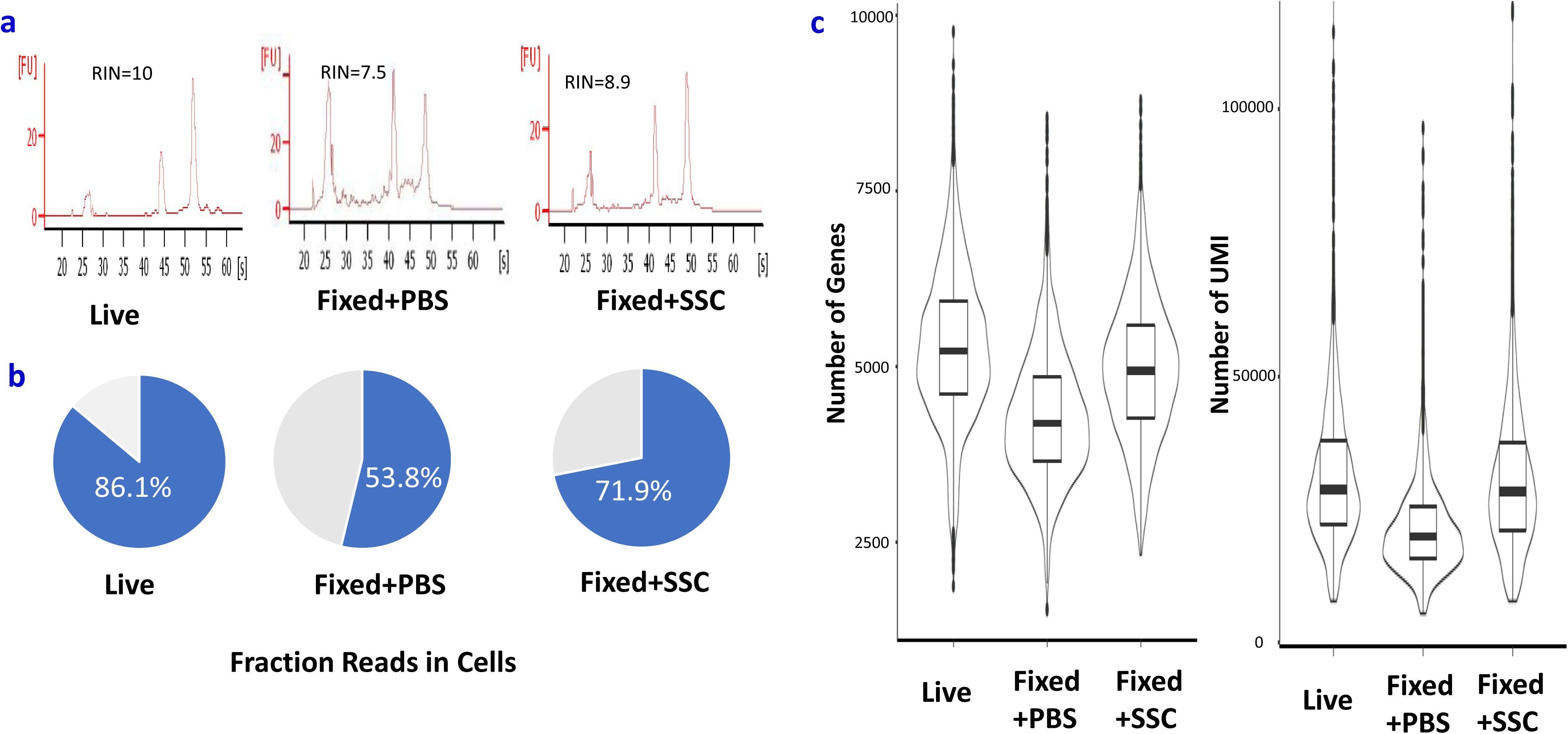
RNA quality and sequencing performance also improved in methanol-fixed KLM1 cells resuspended in SSC. **a**. RNA was partially degraded in fixed KLM1 cells rehydrated in PBS. Cells resuspended in SSC kept RNA high quality. **b**. The genes and UMI counts detected dropped about 20% and 30% respectively in PBS-rehydrated KLM1 cells while almost no drop in SSC-resuspended cells. **c**. “fraction reads in cells” in PBS-rehydrated KLM1 cells was very low, indicating leakage of RNA from cells. In contrast, KLM1 samples resuspended in SSC had much higher “fraction reads in cells”, meaning it prevented the cytoplasmic RNA leakage after the fixative was removed.

### Distinct subpopulations could be detected in fixed PBMCs

To characterize cellular heterogeneity among fixed PBMCs, PCA was run on the top 1000 variable genes ranked by their normalized dispersion as described by Zheng et al [7]. Graph-based clustering identified nine distinct cell clusters in one fixed sample from donor DTM-Y (Figure 5b), which were visualized in two-dimensional projection of t-SNE. To identify cluster-specific genes, differential expression of each gene was calculated between that cluster and the average of the rest of clusters. Some well-known markers of immune cell types (Table S2) were detected in 3 W fixed cell clusters and were used for classification of PBMCs (Figure 5c). Examination of these cluster-specific genes revealed major subpopulations of PBMCs at expected ratios (StemCell Technologies, Document #23629): ~ 55% T cells (enrichment of CD3D in clusters 2, 3, 4, 6 and 8), ~ 7% NK cells (enrichment of NKG7 in cluster 9), ~ 9% B cells (enrichment of CD79A in cluster 7) and ~ 27% myeloid cells (enrichment of S100A8 in clusters 1and 5). Finer substructures were detected with the T-cell cluster: clusters 2, 3, 4 and 8 were CD4+ T cells (IL7 R-enriched), whereas cluster 6 was CD8+ T cells (CD8A-enriched). However, the boundaries among CD4+ and CD8+ and NK cells were blurred. This observation is in agreement with the report from Zheng et al [7]. To identify subpopulations within the myeloid population, k-means clustering was further applied in cluster 1 and 5. Three subtypes were found: CD14+ monocyte (CD14-enriched), CD16+ monocyte (FCGR3A-enriched) and dendritic cells (FCER1A-enriched). Overall, the above results demonstrated that all major subtypes could be detected in our fixed PBMC sample using scRNA-Seq. When we looked at the tSNE projection, we did find the changes of the relative distances of the clusters due to the loss of genes detected after fixation (Figure S3). However, the low abundant populations (B, NK, DC) in each sample were still detected. Furthermore, subpopulations were detected from fixed PBMCs at a similar proportion to those of live PBMCs (Table 1), demonstrating fixation did not impact the resolution of detected population.

**Figure 5.**
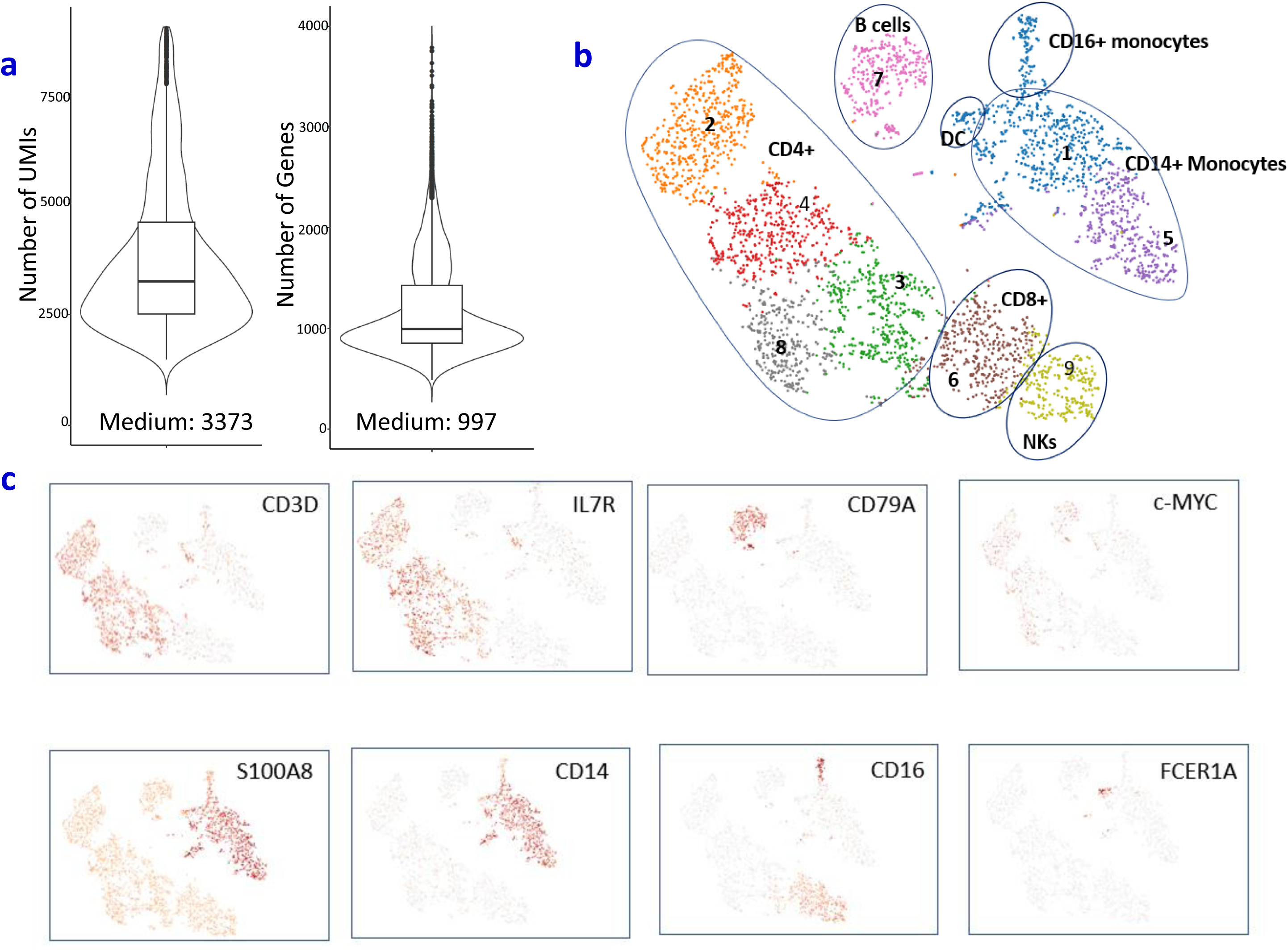
Major PBMC subtypes can be identified in fixed sample. **a**. Distribution and the medium numbers of genes and UMI detected per cell in fixed PBMCs from donor DTM-Y. **b**. tSNE projection of the same fixed PBMCs. Cells were grouped into nine clusters. Classification of PBMCs was inferred from the annotation of cluster-specific genes, and based on expression of some well-known markers of immune cell types. Furthermore, subpopulations were detected from fixed PBMCs at a similar proportion to those of live PBMCs from Donor DTM-X (Table 1). **c**. Some marker genes can be detected in fixed sample. In addition, c-Myc gene was also detected in many cells after fixation although it is quite unstable and easily degraded.

**Table 1.** Proportion of major cell types detected in live and fixed PBMC

| Donor ID | PBMC Processing | Number of Cells Captured | Myeloid Cells |  | Lymphoid Cells |  |  |
| --- | --- | --- | --- | --- | --- | --- | --- |
|  |  |  | Dendritic Cells | Monocytes | B Cells | T Cells | NK Cells |
| DTM-X | Live+PBS | 4,975 | 105 (2.11%) | 1442 (28.98%) | 230 (4.62%) | 2774 (55.76%) | 365 (7.34%) |
| DTM-X | Methanol-3H+SSC | 2,933 | 56 (1.91%) | 818 (27.89%) | 175 (5.97%) | 1622 (55.30%) | 242 (8.25%) |
| DTM-X | Methanol-3W+SSC | 2,826 | 50 (1.77%) | 783 (27.71%) | 196 (6.97%) | 1513 (53.54%) | 228 (8.07%) |
| DTM-Y | Live+PBS | 5,883 | 141 (2.40%) | 1815 (30.85%) | 473 (8.04%) | 2900 (49.29%) | 354 (6.02%) |
| DTM-Y | Methanol-3W+SSC | 3,899 | 64 (1.64%) | 982 (25.19%) | 358 (9.18%) | 2165 (55.53%) | 272 (6.98%) |

It is well known that c-myc is an unstable mRNA [18] in many cells. We detected its expression in many fixed PBMCs (Figure 4b), further demonstrating that our fixation procedure is an efficient method for stabilization of RNA which undergoes rapid changes.

## Discussion

There is a high demand for methods that allow disconnecting time and location of sampling from subsequent single-cell analysis. Here we, for the first time, present a new methanol-fixation processing procedure that prevents PBMC RNA from degradation and loss and is compatible with 10x Chromium standard droplet-base scRNA-Seq. We demonstrate that fixation of PBMCs did not alter their transcriptional profiles and gene expression levels. The protocol was confirmed with CD8+ T cell scRNA-Seq. It also improved the scRNA-Seq performance in three primary cell types and three cell lines. This fixation and resuspension method remains an accurate, sensitive, reproducible and comprehensive characterization of RNAs in a single cell.

Fixation is a process that helps to lock nucleic acids and proteins in place within cells. Unlike aldehydes, alcohol fixatives remove and replace free water and cause a change in the tertiary structure of nucleic acids and proteins by destabilizing hydrophobic bonding, but do not covalently modify them. After alcohol fixation, cells are placed from an aqueous environment to a non-aqueous environment. Alcohols disable intrinsic biomolecules—particularly proteases and RNases —which otherwise digest or damage the sample RNA. However, alcohols do not inactivate RNase completely. After alcohol removal, endogenous RNase may be reactivated during the rehydration of cells in PBS (Figure 1b and S1b). RNA hydrolysis may be of little importance for real time qPCR analysis of short amplicons in some cell lines [19], but it hampers the analysis of complete full-length RNA molecules in droplet-based scRNA-Seq. In our protocol, fixed PBMCs were resuspended in 3X or 5X SSC buffer, a high salt solution. High salt buffer also denatures the protein and nucleic acid and is reported to improve RNA quality in fixed and permeabilized cells [16]. Therefore cells resuspended in high concentration of SSC buffer did not rehydrate. That may explain why this processing preserved the RNA integrity. After cell suspension is added to reverse transcription master mix, the SSC is diluted. It is critical to quickly load the mixed suspension to chip and instrument for GEM generation and RT reaction.

A key challenge across all single-cell RNA-sequencing (scRNA-Seq) techniques is the preservation of each cell’s transcriptional profile throughout the entire sample handling process. The ideal preservation protocol is to prevent or arrest the degenerative processes as soon as a tissue or cell is deprived of its blood supply. In this way autolysis is inhibited and loss and diffusion of soluble substances can be avoided during tissue dissociation or cell sorting. However, cell fixation is often carried out after, not before, single cell preparation [9, 10, 14] because the fixative may interfere with enzyme digestion or antibody binding to the cells. It should be noted that SSC may not be an issue. Nilsson et al demonstrate that antibody staining and FACS are not compromised in the presence of 2 M or 4 M NaCl [16]. 3X SSC is composed of 0.45 M NaCl and 0.045 M trisodium citrate. SSC should not have a major impact on antibody binding either although this is not tested in this study. SSC is a buffer commonly used in hybridization solution. It is reported that 5 x SSC could be used as sheath fluid without disturbing cell-cycle flow analysis [18]. It does not affect the staining and sorting of fixed cells which have been stained with the specific DNA fluorochrome Hoechst 33258. This allows for cells to be fixed right after dissociation and before slow flow cell sorting. Unfortunately this fixation method does not work for whole blood, in particular it is problematic for neutrophils (unpublished data). We found that even live neutrophils did not work with 10 X Chromium single-cell system probably because this complex cell type is easy to activate and undergo rapid RNA damage.

Another reported preservation method is cryopreservation followed by resuscitation for subsequent processing [20]. Although frozen samples are compatible with Droplet-based sequencing [7], it remains to be determined what happens to cells with freezing and thawing manipulation and in unfavorable temperature and medium conditions. Freezing medium components such as DMSO are toxic to cells and may influence gene expression. In Guillaumet-Adkins et al’s report, the freezing process resulted in as high as 23% damaged cells, evidenced by the positive staining with propidium iodine [20]. In contrast, in our study low quality cells did not elevate after methanol-fixation processing (Figure 2b). Of course during cell fixation and the steps that follow there are also substantial changes to the composition and appearance of cell and tissue components, and these are quite far removed from the ideal “life-like state”. However, our data and previous reports [9] showed the transcriptome profiles and gene expression levels were well preserved after fixation. This conservation allows a dynamic ever-changing intracellular environment “fixed” at a given cellular state. Technically the cryopreservation protocol has several cycles of pelleting and washing and usually requires a large number of starting material, i.e. millions of cells. In our study, the CD8+ T cell number was only 0.06 million and could suffer from high speed centrifugation which could result in cell loss. Another advantage of fixation over cryopreservation is convenience. Fixation protocol is much simpler and faster than cryopreservation. It does not require a liquid nitrogen freezer for cell preservation and transportation. In addition, the use of fixed cells eliminates all the problems associated with the manipulation of fresh cells.

## Conclusions

The developed fixation protocol is simple and convenient and has little impact on single cell transcriptome profiles. It would be suitable for scRNA-Seq analysis of many primary tissues with a high content in proteases and RNases such as pancreas, skin or lymphatic and immune tissues. It also could lead to a paradigm shift for complex single-cell study design when standardization (such as magnet stirring) is developed and implemented.

## Abbreviations

DTT: dithiothreitol; GEM: gel bead-in-emulsion; PBMC: peripheral blood mononuclear cells; PCA: principal component analysis; RIN: RNA integrity number; SSC: saline sodium citrate; scRNA-Seq: single-cell RNA sequencing; SMART: switching mechanism at 5’ end of RNA template; t-SNE: t-distributed Stochastic Neighbor Embedding; UMI: Unique molecular identifier

## Acknowledgements

We thank Andrew Martin for discussions and comments on the manuscript.

## Authors’ contribution

JC conceived the study; JST defined the broader strategy and supervised. JC, RS and HZ performed the experiments from sample prep, to library generation to next-generation sequencing with help from WL. FC developed the single-cell RNA sequencing processing pipelines and performed the statistical analysis with support of JC, JMC, YK and JST. JC wrote the manuscript and prepared the figures. JST and KRS discussed and edited the manuscript. All authors read and approved the final manuscript.

## Consent for publication

Not applicable.

## Ethics approval and consent to participate

Not applicable.

## Data availability

Raw sequencing data in BAM format as well as filtered gene-barcode matrices have been deposited at NCBI Gene Expression Omnibus (GEO) and are accessible through accession number GSE112845.

---

**Table 1.** Proportion of major cell types detected in live and fixed PBMC

**Additional File 1: Table S1.** Potential impacts on single cell transcriptome analysis after cell fixation

**Additional File 2: Table S2.** Marker genes used for identifying PBMC subpopulations

**Additional File 3: Figures S1.** Resuspension in 3X SSC preserved cell RNA integrity. **a**. The methanol-fixed PBMCs resuspended in 3X or 5X SSC buffer showed high quality of RNA determined with Bioanalyzer traces. **b**. The new processing method was validated in several other cell types resuspended in 3X SSC for 30 min. The RIN numbers were significantly improved in primary cell types and cell lines with a p value of 0.00001 and 0.008 respectively.

**Additional File 4: Figure S2.** Fixation processing did not aggregate the cells. PBMCs with greater than 2500 genes detected (**a**) or CD8+ cells with more than 2000 genes detected (**b**) were flagged because these droplets usually contain more than one cell (doublet or multiplet). Fixation did not increase the rate of these GEMs/cells. Methanol-fixed PBMCs remained microscopically visible as single, intact round cells with the similar size as live ones.

**Additional File 5: Table S3.** Sequencing metrics summary of ten scRNA-Seq datasets

**Additional File 6: Figure S3.** tSNE projection of live and fixed PBMCs from donor DTM-X (**a**) and donor (**b**). Cells were grouped using graph-based method. Classification of PBMCs was inferred from the annotation of cluster-specific genes, and based on expression of some well-known markers of immune cell types. Although fixation lead to changes of the relative distances of the clusters due to the loss of genes detected, it did not impact the resolution of the low abundant populations (B, NK, DC) in each sample. Subpopulations were detected from fixed PBMCs at a similar proportion to those of live PBMCs (Table 1).

## References

1. Proserpio V, Mahata B: Single-cell technologies to study the immune system. Immunology 2016, 147(2):133–140.

2. Buchholz VR, Flossdorf M: Single-Cell Resolution of T Cell Immune Responses. In: Advances in Immunology. Academic Press; 2018.

3. Grun D, van Oudenaarden A: Design and Analysis of Single-Cell Sequencing Experiments. Cell 2015, 163(4):799–810.

4. Bacher R, Kendziorski C: Design and computational analysis of single-cell RNA-sequencing experiments. Genome Biol 2016, 17:63.

5. Wu AR, Neff NF, Kalisky T, Dalerba P, Treutlein B, Rothenberg ME, Mburu FM, Mantalas GL, Sim S, Clarke MF et al: Quantitative assessment of single-cell RNA-sequencing methods. Nat Methods 2014, 11(1):41–46.

6. Macosko EZ, Basu A, Satija R, Nemesh J, Shekhar K, Goldman M, Tirosh I, Bialas AR, Kamitaki N, Martersteck EM et al: Highly Parallel Genome-wide Expression Profiling of Individual Cells Using Nanoliter Droplets. Cell 2015, 161(5):1202–1214.

7. Zheng GX, Terry JM, Belgrader P, Ryvkin P, Bent ZW, Wilson R, Ziraldo SB, Wheeler TD, McDermott GP, Zhu J et al: Massively parallel digital transcriptional profiling of single cells. Nat Commun 2017, 8:14049.

8. Hicks SC, Townes FW, Teng M, Irizarry RA: Missing data and technical variability in single-cell RNA-sequencing experiments. Biostatistics 2017.

9. Alles J, Karaiskos N, Praktiknjo SD, Grosswendt S, Wahle P, Ruffault PL, Ayoub S, Schreyer L, Boltengagen A, Birchmeier C et al: Cell fixation and preservation for droplet-based single-cell transcriptomics. BMC Biol 2017, 15(1):44.

10. Cao J, Packer JS, Ramani V, Cusanovich DA, Huynh C, Daza R, Qiu X, Lee C, Furlan SN, Steemers FJ et al: Comprehensive single-cell transcriptional profiling of a multicellular organism. Science 2017, 357(6352):661–667.

11. Esser C, Gottlinger C, Kremer J, Hundeiker C, Radbruch A: Isolation of full-size mRNA from ethanol-fixed cells after cellular immunofluorescence staining and fluorescence-activated cell sorting (FACS). Cytometry 1995, 21(4):382–386.

12. Ilicic T, Kim JK, Kolodziejczyk AA, Bagger FO, McCarthy DJ, Marioni JC, Teichmann SA: Classification of low quality cells from single-cell RNA-seq data. Genome Biol 2016, 17:29.

13. Yamada H, Maruo R, Watanabe M, Hidaka Y, Iwatani Y, Takano T: Messenger RNA quantification after fluorescence activated cell sorting using intracellular antigens. Biochem Biophys Res Commun 2010, 397(3):425–428.

14. Karaiskos N, Wahle P, Alles J, Boltengagen A, Ayoub S, Kipar C, Kocks C, Rajewsky N, Zinzen RP: The Drosophila embryo at single-cell transcriptome resolution. Science 2017, 358(6360):194–199.

15. Kozubek S, Lukasova E, Amrichova J, Kozubek M, Liskova A, Slotova J: Influence of cell fixation on chromatin topography. Anal Biochem 2000, 282(1):29–38.

16. Nilsson H, Krawczyk KM, Johansson ME: High salt buffer improves integrity of RNA after fluorescence-activated cell sorting of intracellular labeled cells. J Biotechnol 2014, 192 Pt A:62–65.

17. Maeda T, Date A, Watanabe M, Hidaka Y, Iwatani Y, Takano T: Optimization of Recovery and Analysis of RNA in Sorted Cells in mRNA Quantification After Fluorescence-activated Cell Sorting. Ann Clin Lab Sci 2016, 46(6):571–577.

18. Khochbin S, Grunwald D, Pabion M, Lawrence JJ: Recovery of RNA from flow-sorted fixed cells. Cytometry 1990, 11(8):869–874.

19. Date A, Maeda T, Watanabe M, Hidaka Y, Iwatani Y, Takano T: An improved protocol for mRNA quantification after fluorescence-activated cell sorting with an increased signal to noise ratio in flow cytometry. Mol Biotechnol 2014, 56(7):591–598.

20. Guillaumet-Adkins A, Rodriguez-Esteban G, Mereu E, Mendez-Lago M, Jaitin DA, Villanueva A, Vidal A, Martinez-Marti A, Felip E, Vivancos A et al: Single-cell transcriptome conservation in cryopreserved cells and tissues. Genome Biol 2017, 18(1):45.

